# Dosing and Serostatus Shape the Efficacy of Adenovirus, mRNA, and Protein Vaccines

**DOI:** 10.1101/2025.07.16.665159

**Authors:** Bakare Awakoaiye, Shiyi Li, Sarah Sanchez, Tanushree Dangi, Nahid Irani, Laura Arroyo, Gabriel Arellano, Shadi Mohammadabadi, Malika Aid, Pablo Penaloza-MacMaster

## Abstract

Despite the widespread use of adenovirus, mRNA, and protein-based vaccines during the COVID-19 pandemic, their relative immunological profiles and protective efficacies remain incompletely defined. Here, we compared antigen kinetics, innate and adaptive immune responses, and protective efficacy following Ad5, mRNA, and protein vaccination in mice. Ad5 induced the most sustained antigen expression, but mRNA induced the most potent interferon responses, associated with robust antigen presentation and costimulation. Unlike Ad5 vaccines, which were hindered by pre-existing vector immunity, mRNA vaccines retained efficacy after repeated use. As a single-dose regimen, Ad5 vaccines elicited superior immune responses. However, as a prime-boost regimen, and particularly in Ad5 seropositive mice, mRNA vaccines outperformed the other vaccine platforms. These findings highlight strengths of each vaccine platform and underscore the importance of host serostatus in determining optimal vaccine performance.

## INTRODUCTION

The rapid development of vaccines based on different platforms, including adenoviral vector vaccines, mRNA vaccines, and protein subunit vaccines, was pivotal in curbing severe disease and death during the COVID-19 pandemic. Early in the pandemic, Ad5-based vaccines such as CanSino and Sputnik V, alongside mRNA vaccines developed by Pfizer-BioNTech and Moderna, provided efficacy against severe SARS-CoV-infection. Protein-based vaccines like Novavax were introduced later but also demonstrated efficacy. While each of these platforms has proven immunogenic and protective in clinical studies, their relative immunologic profiles and efficacy have not been directly compared in controlled settings.

Among COVID-19 vaccines, Ad5-based and mRNA-based vaccines emerged as leading platforms due to their potent immunogenicity. However, the influence of vaccine schedule and host serological status remains underexplored. Adenoviral vectors like Ad5 are known to elicit strong and durable T cell responses, yet their efficacy can be reduced in individuals with pre-existing immunity to the vector, particularly in populations with high adenovirus seroprevalence^1^. In this study, we performed a head-to-head comparison of Ad5, mRNA, and protein vaccines in mice to examine antigen expression dynamics, innate and adaptive immune responses, and protective efficacy. Our results reveal that the performance of each platform is context-dependent, shaped by the vaccination schedule and the host’s serostatus. These findings provide immunological insights into three widely used vaccine platforms and offer guidance on tailoring vaccine strategies to specific populations.

## RESULTS

### Antigen presentation, costimulation, and cytokine expression after vaccination with Ad5, mRNA, and protein

There are three critical signals necessary for the activation of adaptive immune responses following natural infection or vaccination: (1) antigen presentation, (2) costimulation, and (3) cytokine signaling ^2^. We conducted experiments to compare these three signals after vaccination with either Ad5, mRNA, or protein (administered 1:10 with AdjuPhos). To examine antigen levels, we vaccinated BALB/c mice intramuscularly with Ad5 or mRNA expressing a luciferase reporter (Ad5-Luc or mRNA-Luc), or luciferase protein itself, and then injected these mice with luciferin at various time points to quantify antigen levels by in vivo bioluminescence (**Fig. 1A**). We utilized BALB/c mice because their white coat facilitates the visualization of the luciferase reporter at the site of vaccination, the quadriceps muscle. mRNA vaccination resulted in diffuse antigen expression extending beyond the injection site, while antigen expression from Ad5 and protein vaccines remained localized at a more discrete site within the quadriceps (**Fig. 1B**). mRNA vaccination induced strikingly high antigen levels at 6 h, but antigen levels rapidly declined after 24 h (**Fig. 1B-1C**). In contrast, the Ad5 vaccine induced low antigen levels at 6 h, but antigen expression persisted longer than with the mRNA vaccine (**Fig. 1B-1C**). With the protein vaccine, antigen levels were lower than the other vaccine platforms (**Fig. 1B-1C**).

**Figure 1.**
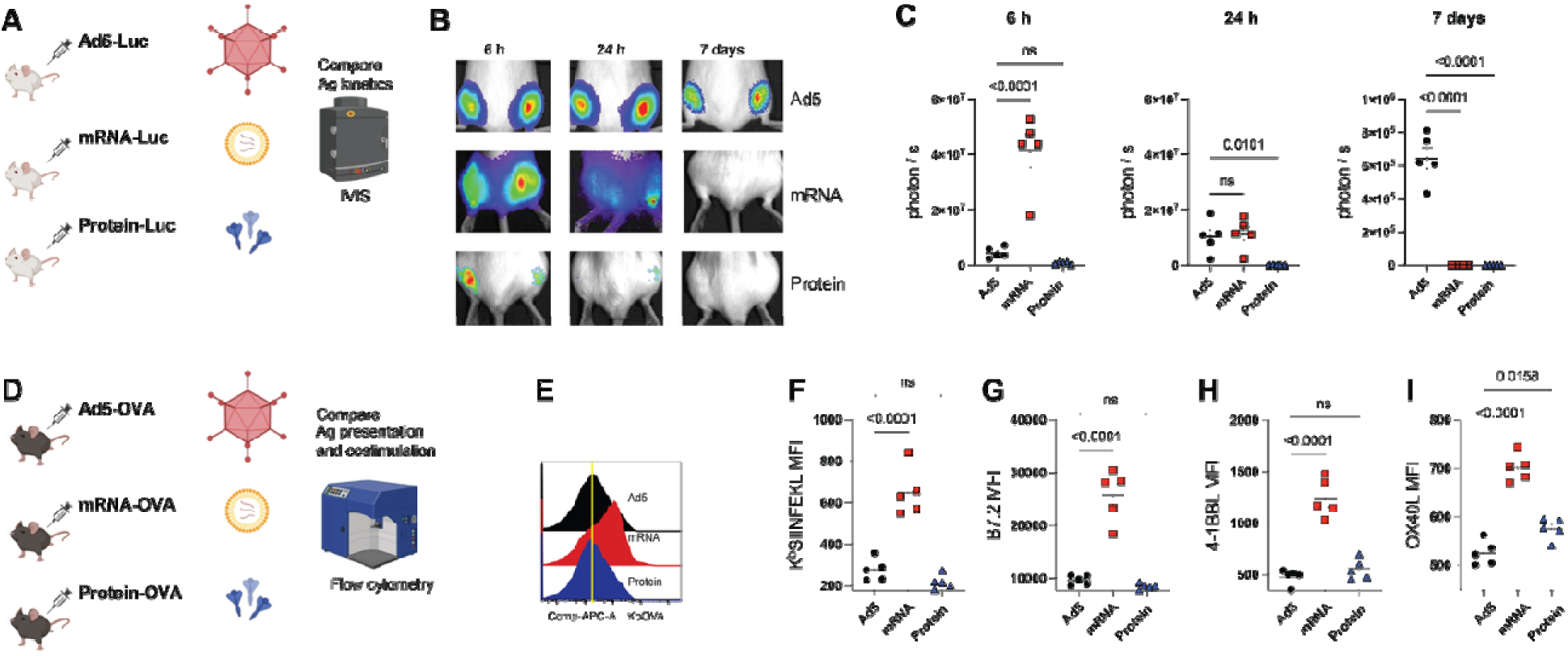
Comparison of antigen presentation and costimulation following immunization with Ad5, mRNA, and protein vaccines. (**A**) Experimental outline for comparing antigen kinetics. BALB/c mice were immunized intramuscularly with Ad5-Luc, mRNA-Luc, or Luciferase. (**B**) In vivo bioluminescence images at various timepoints after immunization. (**C**) Summary of antigen expression by in vivo bioluminescence. (**D**) Experimental outline for interrogating antigen presentation and costimulation at day 1 post-vaccination. (**E**) Representative FACS histograms showing DC that present cognate antigen. (**F**) K^b^SIINFEKL expression. (**G**) B7.2 expression. (**H**) 4-1BBL expression. (**I**) OX40L expression. Data are from one experiment (n=5 per group). Experiments were repeated once with similar results. Indicated *P* values were calculated by ordinary one-way ANOVA with Dunnett’s multiple comparisons. Error bars represent SEM.

We then compared antigen presentation (Signal 1) by dendritic cells (DC) after immunization with the three vaccine platforms (**Fig. 1D**). C57BL/6 mice were vaccinated intramuscularly with Ad5 or mRNA vaccines expressing ovalbumin (Ad5-OVA and mRNA-OVA), or OVA protein itself. At day 3 post-vaccination, we harvested draining lymph nodes and quantified antigen presentation on DCs utilizing a fluorochrome-labeled antibody recognizing MHC class I bound to SIINFEKL peptide (**Fig. 1D**). Interestingly, mRNA vaccination generated DCs with superior levels of MHC class I bound to SIINFEKL peptide, suggesting more robust antigen presentation capacity (**Fig. 1E-1F**). We also compared costimulation (Signal 2) by DC after immunization with the three vaccine platforms. mRNA vaccination induced higher expression of B7.2, 4-1BBL, and OX40L molecules on DCs, relative to the other vaccine platforms, suggesting greater costimulatory potential (**Fig. 1G-1I**). We then compared cytokine responses (Signal 3) at 6 h post-vaccination by Luminex (**Fig. 2A**). mRNA vaccination induced higher levels of various acute cytokines, including interferon related cytokines like gamma-induced protein 10 (IP-10), IFNβ, and IFNγ, relative to the other vaccine platforms (**Fig. 2B**). mRNA vaccination also induced a significant increase in IL-6, but other cytokines like IL-12, TNFα, and IL-1β were comparable across the different vaccine platforms (**Fig. 2C**). Together, these data showed striking differences in antigen presentation, costimulation, and acute cytokine responses between the three vaccine platforms.

**Figure 2.**
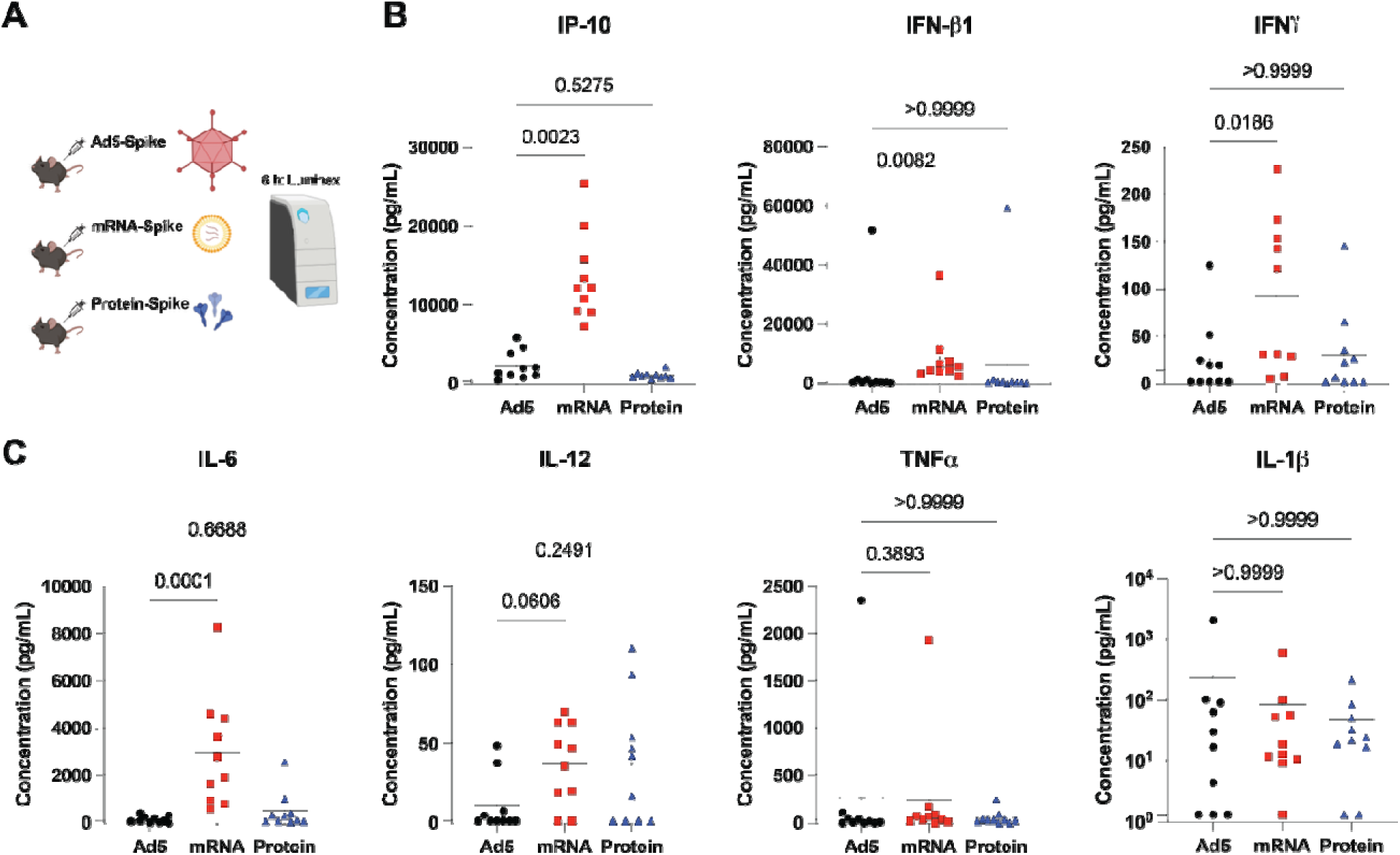
Acute cytokine profiles following immunization with Ad5, mRNA, and protein vaccines. (**A**) Experimental outline for comparing acute cytokine responses at 6 h post-immunization in C57BL/6 mice that were immunized intramuscularly with Ad5, mRNA, and protein vaccines. (**B**) Interferon related cytokines. (**C**) Other cytokines. Data are from two experiments (n=5 per group/experiment). Indicated *P* values were calculated by Kruskal-Walli one-way ANOVA test with Dunn’s multiple comparisons. Error bars represent SEM.

### Single-cell gene expression analyses reveal differences in viral sensing pathways and interferon related pathways

We then performed gene expression analyses after vaccination with Ad5, mRNA, and protein. We harvested draining lymph nodes at day 1 post-vaccination, followed by enrichment of live CD45+ cells by magnetic cell sorting and single-cell RNA sequencing (scRNA-Seq) (**Fig. 3A**). mRNA vaccination induced the most pronounced transcriptional changes with 529 unique genes relative to naïve (**Fig. 3B**). Protein vaccination, on the other hand, induced the least transcriptional changes with only 8 unique genes relative to naïve (**Fig. 3B and Supplemental Table 1**).

**Figure 3.**
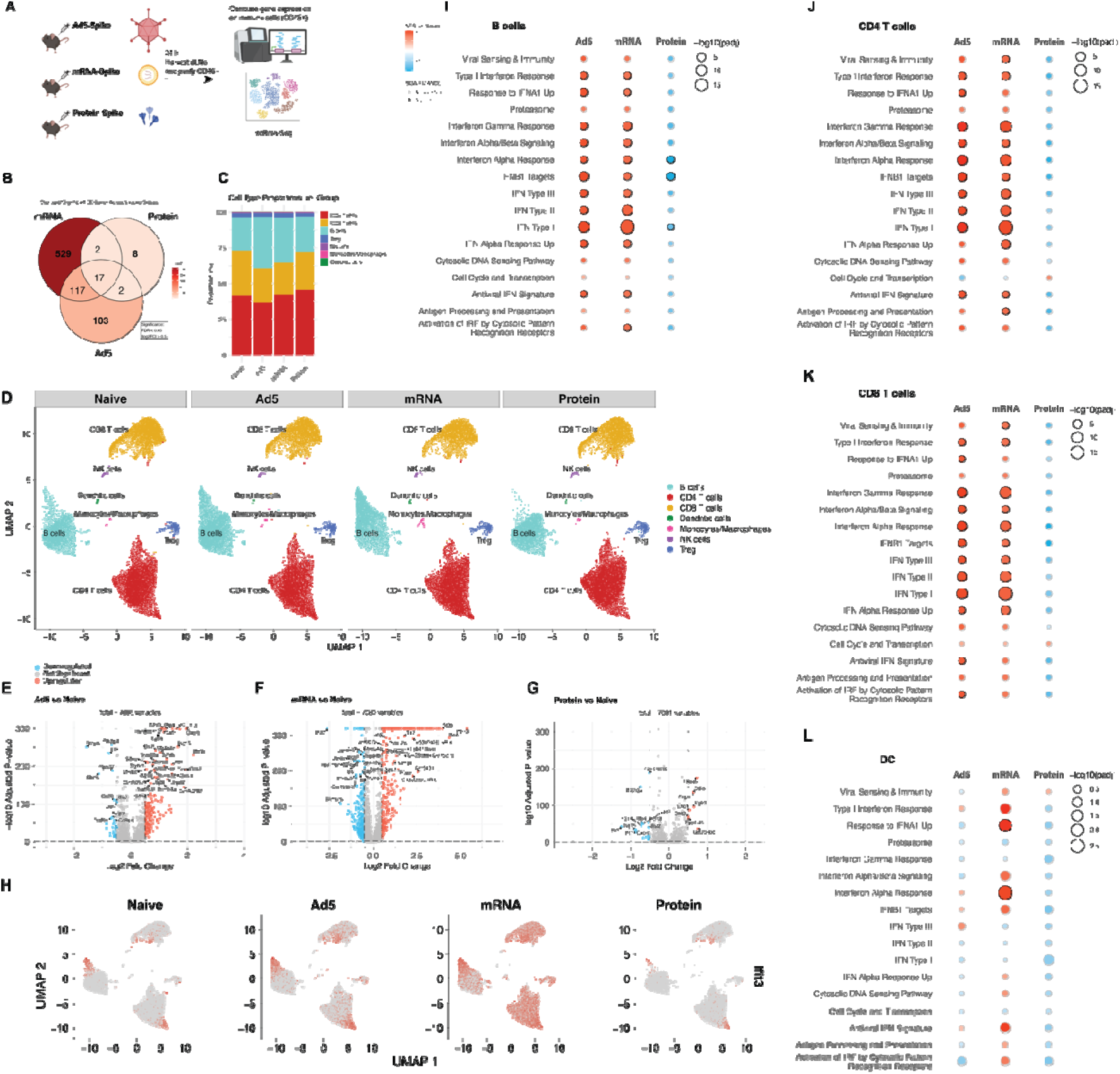
Single cell RNA-Seq (scRNA-Seq) analyses reveal enrichment in interferon pathways and viral sensing pathways in mice vaccinated with mRNA. **(A)** Experimental outline for gene expression analysis of immune cells (CD45+) from draining lymph nodes. C57BL/6 mice were immunized intramuscularly with each respective vaccine, and at 24 h post-immunization, draining lymph nodes were enriched for CD45+ cells using magnetic beads. (**B**) Venn diagram showing the overlap of significant differentially expressed genes (DEGs) among each vaccine group compared to naive controls. Significance was defined as FDR < 0.05 and absolute log2 fold change > 0.5. (**C**) Bar plot depicting the proportions of annotated cell types in each vaccination group. (**D**) UMAP visualization of single-cell transcriptomes, with cells colored and labeled by cell identity. UMAPs are shown separately for each vaccination group. (**E-G**). Volcano plots displaying DEGs for Ad5 vs. naive, mRNA vs. naive, and protein vs. naive comparisons. Vertical lines indicate log2 fold change of 0.5; horizontal lines indicate FDR of 0.05. Genes upregulated in the vaccine group are shown in red, downregulated in blue, and non-significant genes in grey. (**H**) UMAP plots showing the expression of Ifit3 across each vaccination group and naive mice. (**I–L**) Dot plots of enriched pathways identified by GSEA in B cells (**I**), CD4 T cells (**J**), CD8 T cells (**K**), and dendritic cells (**L**) for each vaccine group compared to naive. The y-axis lists pathway names, dot size represents –log10(adjusted p-value), and color corresponds to the normalized enrichment score (NES). A black ring denotes significant enrichment (FDR < 0.05); a grey ring indicates non-significance.

Most cells in the draining lymph node were lymphocytes, with a smaller proportion consisting of monocytes, macrophages, and DC (**Fig. 3C**). Of note, Ad5 and mRNA vaccination were associated with higher frequencies of B cells (**Fig. 3D**). Consistent with the Luminex data that showed higher interferon responses in mRNA vaccinated mice, we observed that mRNA vaccination elicited the most potent expression of IFN-induced genes (ISGs), including those coding for Ifitm3, Ifit3, Ifi44, Ifit3, and Cxcl10 (**Fig. 3E-3H**). We then analyzed inflammatory pathways on specific immune cell subsets, including lymphocytes and DC. When we analyzed lymphocytes, Ad5 and mRNA vaccination were associated with enrichment in pathways involved in viral sensing, IFN-I, IFN-II, IFN-III, proteasome function, antigen processing, antigen presentation, and activation of pattern recognition receptors, relative to protein vaccination (**Fig. 3I-3K**). When we analyzed gene expression on DC specifically, mRNA vaccination was associated with enrichment in pathways involved in viral sensing, IFN-I, and activation of pattern recognition receptors (**Fig. 3L**). These data demonstrate that mRNA vaccination is associated with a more pronounced enrichment in IFN-I induced genes especially on DC, relative to the other vaccine platforms.

### Comparative analyses of adaptive immune responses

We then compared T cell and antibody responses after vaccination. C57BL/6 mice were vaccinated intramuscularly with Ad5 or mRNA vaccines expressing the SARS-CoV-2 spike protein (Ad5-spike and mRNA-spike), or SARS-CoV-2 spike protein itself, and analyzed CD8 T cell responses by tetramer stains and antibody responses by enzyme-linked immunosorbent assays (ELISA) (**Fig. 4A**). After a single prime, Ad5 elicited the most potent CD8 T cell response (**Fig. 4B**). However, following a booster vaccination, CD8 T cell responses were comparable among the Ad5 and mRNA groups. Protein vaccination elicited low CD8 T cell responses that were near the limit of detection. We also compared antibody responses between the different vaccine platforms and observed greater responses with Ad5 vaccines, compared to the other vaccines (**Fig. 4C**). Overall, these initial data showed that Ad5 is more immunogenic than the other vaccines when comparing single dose regimens.

**Figure 4.**
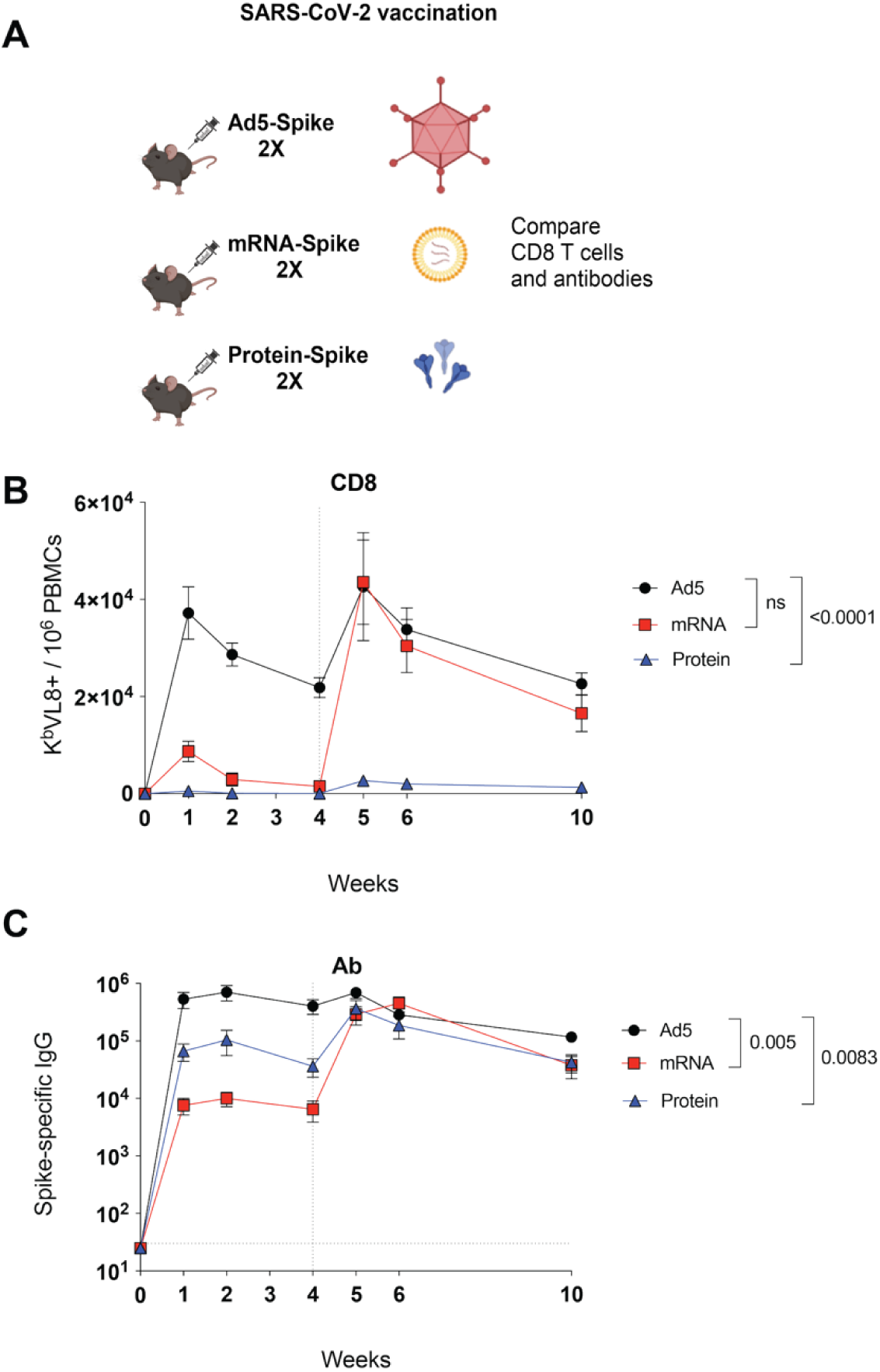
Comparative analyses of adaptive immune responses. (**A**) Experimental outline for comparing antibody and CD8 T cell response following vaccination. C57BL/6 mice were immunized intramuscularly with each respective vaccine based on the SARS-CoV-2 spike antigen. After 4 weeks, mice were boosted homologously. (**B**) Summary of SARS-CoV-2-specific (K^b^ VL8+) CD8^+^ T cells in PBMCs. (**C**) Summary of SARS-CoV-2-specific antibody titers in sera. Data from four experiments (n=5 mice per group/experiment). The vertical dashed line indicates the time of boosting. Indicated *P* values were calculated by ordinary one-way ANOVA with Dunnett’s multiple comparisons at week 10. Error bars represent SEM.

Ad5 is highly seroprevalent in humans ^1^. We performed a retrospective study to measure the titers of Ad5 hexon-specific antibodies using de-identified sera from donors selected across three independent human cohorts within the United States. We observed 100% seropositivity to Ad5, likely reflecting prior infections with this adenovirus, which is known to cause common colds. (**Fig. 5**). This pattern of Ad5 seropositivity was uniformly observed among all human volunteers. Given this widespread seroprevalence, we reasoned that a more appropriate comparison of vaccine platforms would need to account for the high level of Ad5 seropositivity in the human population. To model this, we first rendered mice seropositive to Ad5 by immunizing them three times with an Ad5 vector devoid of any vaccine antigen (Ad5-Empty) (**Fig. 6A**). This regimen generated high titers of Ad5-specific antibodies above >10 endpoint titer, similar to those observed in humans (**Fig. 6B**). These Ad5-seropositive mice were then immunized with vaccines based on Ad5, mRNA, or protein, and adaptive immune responses were evaluated. In Ad5-seropositive mice, a prime-boost regimen with mRNA elicited more robust CD8 T cell responses than the other vaccine regimens (**Fig. 6C–6F**). Moreover, CD8 T cells induced by the mRNA regimen showed higher co-expression of cytokines such as IFNγ, TNFα, and IL-2 (**Fig. S1A– S1C**). We did not observe statistically significant differences in CD4 T cell responses between the different vaccine platforms (**Fig. S1D**).

**Figure 5.**
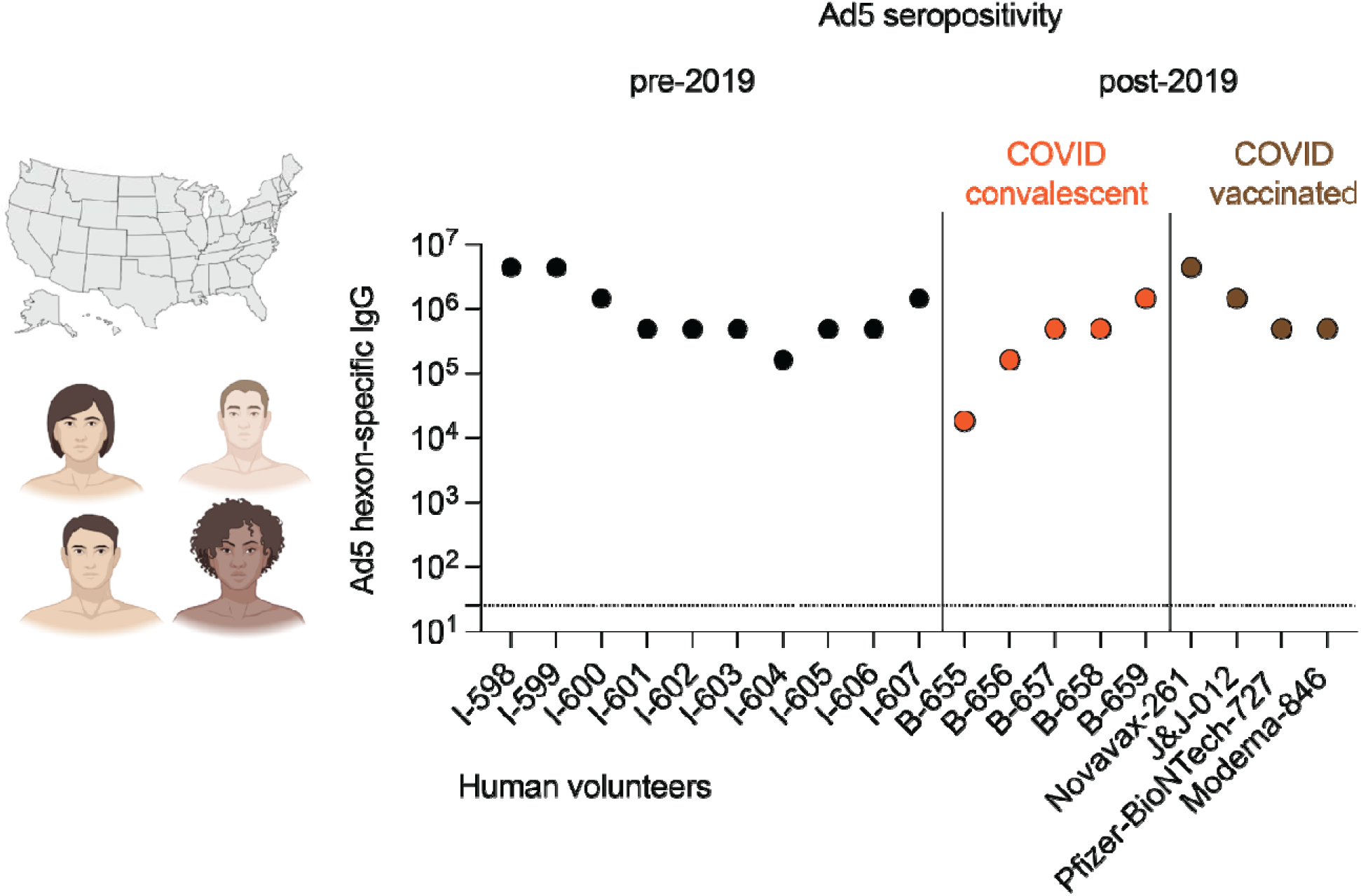
Ad5 seropositivity in human serum samples. Ad5-specific antibody titers in human sera samples from three independent human cohorts. Each data point represents individual human samples. Data are from de-identified human serum samples obtained from other studies (secondary research) and grouped based on whether they were collected before or after 2019. The horizontal dashed line represents LOD.

**Figure 6.**
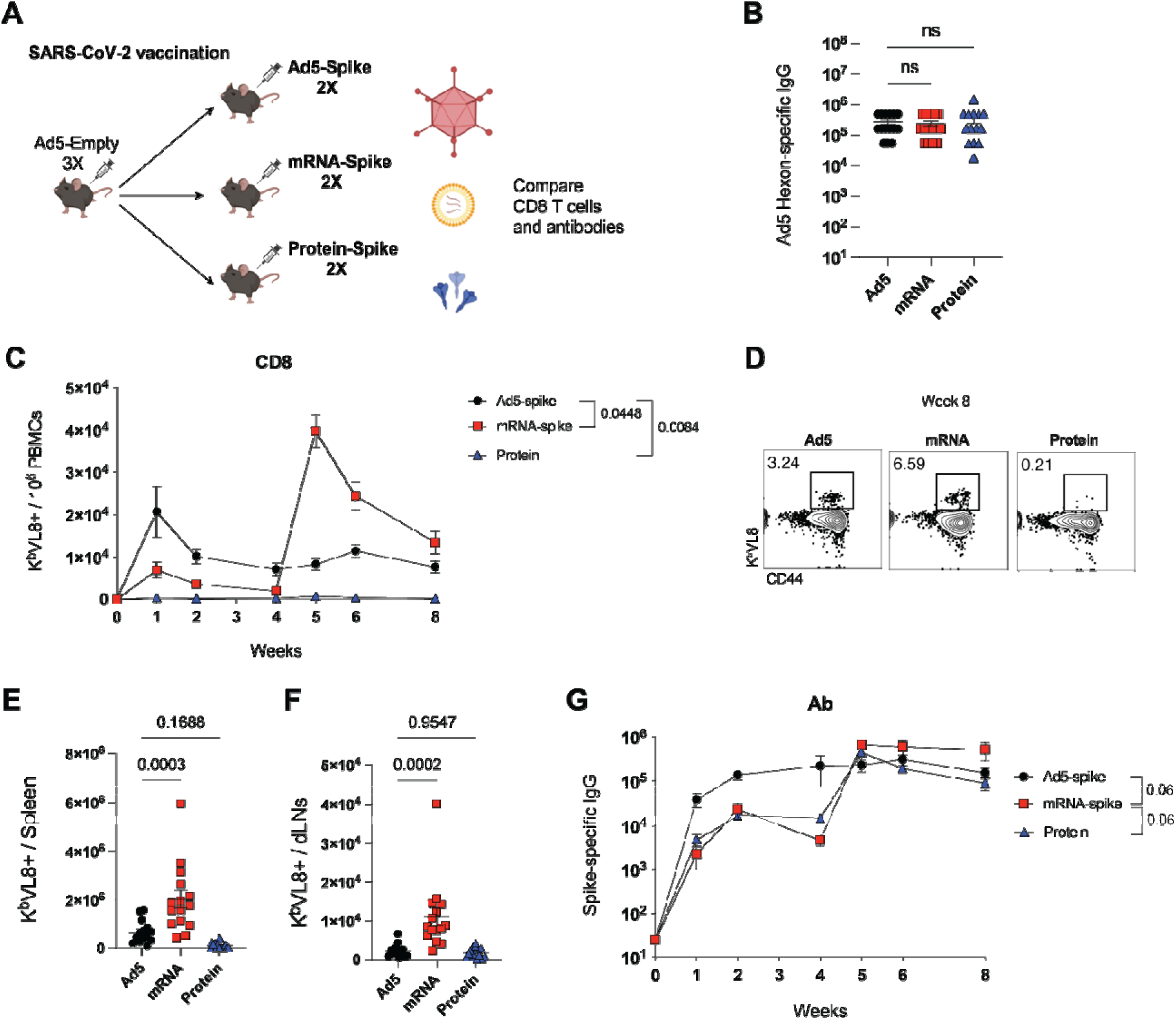
Comparative analyses of adaptive immune responses in Ad5 seropositive hosts. (**A**) Experimental outline for comparing vaccine-elicited responses. First, mice were made Ad5 seropositive by injecting them with 10^9^ PFUs of Ad5-Empty per mouse, once every 3 weeks for a total of three doses. These mice were then immunized intramuscularly with each respective vaccine based on the SARS-CoV-2 spike antigen. After 4 weeks, mice were boosted homologously. (**B**) Ad5 seropositivity was confirmed by Ad5 hexon-specific antibody titers. (**C**) Summary of SARS-CoV-2-specific (K^b^ VL8+) CD8^+^ T cells in PBMCs. (**D**) Representative FACS plots showing SARS-CoV-2-specific CD8^+^ T cell in PBMCs at week 8. (**E**) Summary of SARS-CoV-2-specific CD8^+^ T cells in spleen. (**F**) Summary of SARS-CoV-2-specific CD8^+^ T cells in draining lymph nodes. (**G**) Summary of SARS-CoV-2-specific antibody titers in sera. Data from four experiments (n=5 mice per group/experiment). The vertical dashed line indicates the time of boosting. Indicated *P* values in panels C, E, F were calculated by ordinary one-way ANOVA with Dunnett’s multiple comparisons (p value for panel C is from week 8). Indicated *P* values in panel F were calculated by two-way ANOVA with Holm-Šídák’s multiple comparisons test (p value for panel F is from week 8). Error bars represent SEM.

We also compared CD8 T cell subset differentiation, and we showed that most antigen-specific CD8 T cells generated by Ad5 exhibited a short-lived effector cell (SLEC) phenotype, whereas most antigen-specific CD8 T cells generated by mRNA and protein showed a memory precursor effector cell (MPEC) phenotype (**Fig. S2A-S2C**). There was also a pattern of improved antibody responses after mRNA vaccination, but the difference was not statistically significant (p=0.06) (**Fig. 6G**). Overall, these data demonstrate that in an Ad5 seropositive host, a prime-boost regimen with mRNA is more immunogenic than the other vaccine platforms.

### Effects of pre-existing immunity

As previously discussed, a major limitation of utilizing Ad5 vectors is their high seroprevalence which can limit their clinical efficacy. In an earlier study, we demonstrated that during Ad5 immunization, pre-existing Ad5-specific antibodies can accelerate the clearance of the Ad5 vector at the site of immunization, limiting antigen expression by the vector ^3^. We also observed that prior immunization of mice with an Ad5-Empty vector limits the immunogenicity of the Ad5 vector platform after subsequent immunizations (**Fig. S3A-S3C**), consistent with prior studies ^4^. These negative effects of pre-existing immunity have been the basis for developing novel adenovirus vector platforms with lower seroprevalence, including Ad26 ^5,6^. Besides using a different adenovirus serotype for booster vaccination, we reasoned that boosting with a protein vaccine could help overcome pre-existing immunity in Ad5-primed mice, potentiating adaptive immune responses. Consistent with our hypothesis, we show that a heterologous Ad5/protein regimen can overcome some of the limitations of the homologous Ad5/Ad5 regimen, resulting in more potent humoral responses (**Fig. S3D-S3I**).

So far, we have shown that prior immunization with Ad5 hampers the re-utilization of this vector, consistent with prior reports. But it is unclear whether the same occurs with mRNA vaccines. Prior studies have shown that mRNA vaccines elicit antibodies against the LNP carrier ^7,8^, but whether these hamper mRNA vaccines remain unknown. To examine whether prior immunization with mRNA vaccines hampers the re-utilization of mRNA vaccines, we immunized mice three times with an irrelevant mRNA vaccine encoding the SARS-CoV-2 spike protein or PBS control. After several weeks, these mice were immunized with an mRNA vaccine encoding a different antigen, the lymphocytic choriomeningitis virus (LCMV) glycoprotein (mRNA-LCMV) (**Fig. S4A**). Interestingly, mice that were previously immunized with the mRNA-spike vaccine generated similar LCMV-specific immune responses, relative to control mice (**Fig.S4B–S4D**). We recapitulated these findings in another model, showing that prior immunization with an mRNA-OVA vaccine did not hamper immune responses to a different mRNA vaccine encoding the mouse hepatitis virus spike protein (mRNA-MHV) (**Fig. S4E– S4G**). Collectively, these data suggest that in contrast to Ad5 vaccines, prior immunization with mRNA vaccines does not impair the re-utilization of mRNA vaccines.

### Comparative efficacy of Ad5, mRNA, and protein vaccines following stringent pathogen challenge

In the prior sections, we have compared immune responses following Ad5, mRNA, and protein vaccination. We did not compare vaccine efficacy using SARS-CoV-2 challenges, because SARS-CoV-2 is rapidly cleared in vaccinated animals. Even a single dose with SARS-CoV-2 vaccines confers robust protection in animal models, making it difficult to ascertain differences in immune protection with our vaccine regimens ^9–14^. To detect differences in immune protection, we relied on a more stringent challenge model based on Listeria monocytogenes. First, Ad5-seropositive mice were vaccinated with Ad5-OVA, mRNA-OVA, or OVA protein, followed by immunogenicity and efficacy studies using a stringent Listeria challenge model (**Fig. 7A**). Consistent with our prior results, the mRNA vaccine regimen elicited greater OVA-specific CD8 T cell responses in blood and tissues, compared to the other vaccine platforms (**Fig. 7B-7G**). There were also improved antibody responses in mRNA vaccinated mice, relative to the other vaccine regimens (**Fig. 7H**). Similar to our studies using the spike antigen model, we did not observe differences in CD4 T cell responses specific for OVA (**Fig. S5A-S5B**). Moreover, we did not observe differences in CD4+ FoxP3+ T regulatory cells (Tregs), suggesting that the differences in vaccine responses were not due to this regulatory cell subset (**Fig. S5C-S5D**).

**Figure 7.**
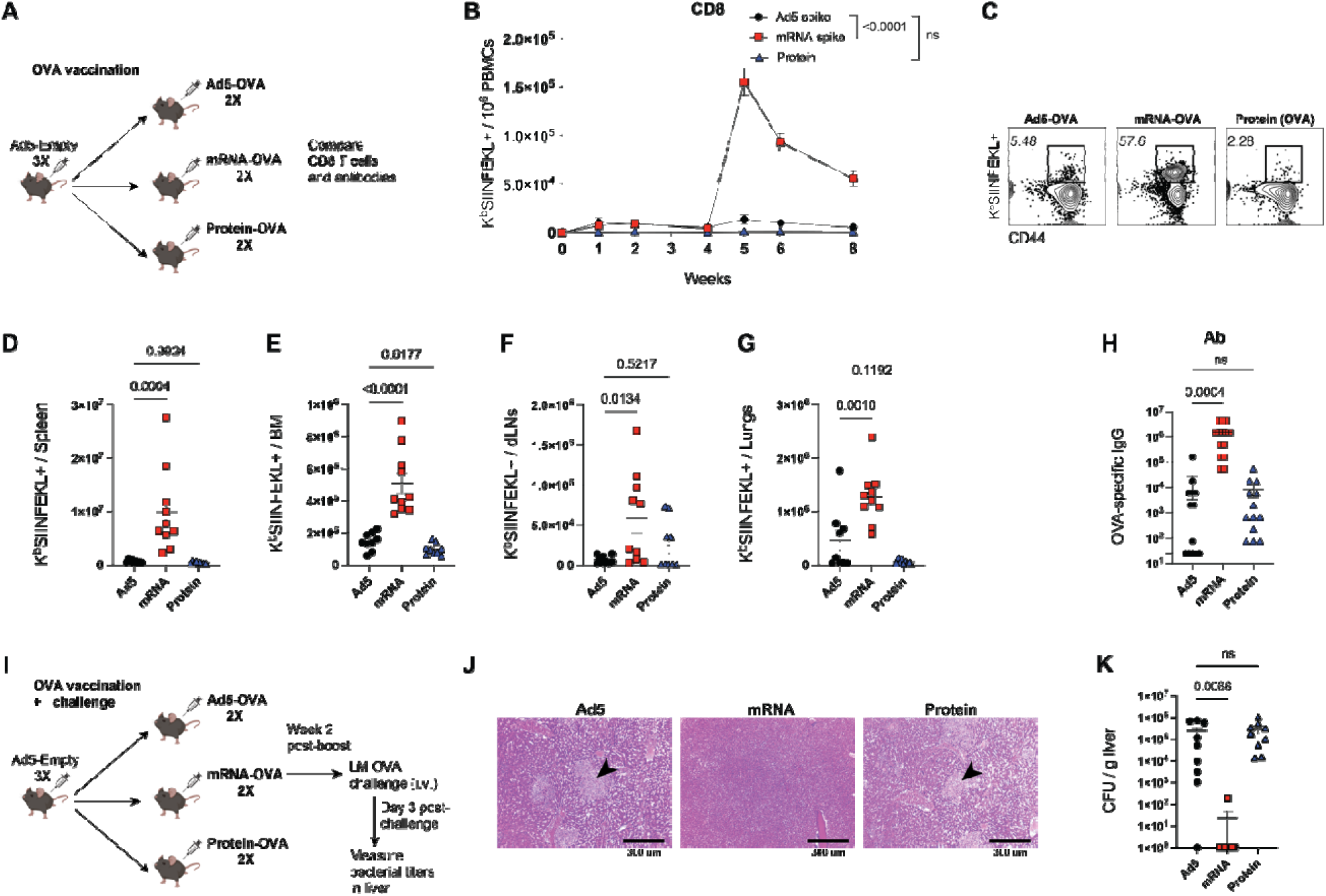
Protective efficacy following vaccination. (**A**) Experimental outline for comparing vaccine-elicited responses and immune protection. First, mice were made Ad5 seropositive by injecting them with 10^9^ PFUs of Ad5-Empty per mouse, once every 3 weeks for a total of three doses. These mice were then immunized intramuscularly with each respective vaccine based on the OVA antigen. After 4 weeks, mice were boosted homologously. (**B**) Summary of OVA-specific (K^b^ SIINFEKL+) CD8^+^ T cells in PBMCs. (**C**) Representative FACS plots showing OVA-specific CD8^+^ T cell in PBMCs. Summary of OVA-specific CD8^+^ T cells in spleen (**D**), bone marrow (**E**), draining lymph nodes (**F**), and lungs (**G**). (**H**) OVA-specific antibody titers in sera. Data from panels C-H are from week ∼8. (**I**) Experimental outline for comparing protective efficacy. On day 3 after LM-OVA challenge, livers were harvested for H&E-stains (**J**) and measuring bacterial loads (**K**). Black arrows in panel J denote areas of necrosis. Data from two experiments (n=4-5 mice per group/experiment). The vertical dashed line indicates the time of boosting. For panels B and D-H, indicated *P* values were determined by ordinary one-way ANOVA test with Dunnett’s multiple comparisons (p values for panels B and H are from week 8). For panel K, indicated *P* values were determined by Kruskal-Wallis test with Dunn’s multiple comparisons. Error bars represent SEM.

To compare vaccine efficacy, we challenged mice intravenously with a supra-lethal dose (10^7^ CFU/mouse) of Listeria monocytogenes expressing OVA (LM-OVA) (**Fig. 7I**). This lethal challenge model is particularly stringent, as most experimental Listeria vaccines are unable to control systemic bacterial dissemination ^15^, rendering it well-suited to compare differences in vaccine efficacy. As expected, mice vaccinated with Ad5 or protein exhibited disseminated necrosis and high bacterial titers in the liver after an LM-OVA challenge (**Fig. 7J-7K**). In contrast, most mice vaccinated with mRNA showed sterilizing immunity, with no evidence of Listeria infection (**Fig. 7J-7K**). These data suggest that mRNA vaccines are more effective than Ad5 vaccines and protein vaccines, especially after a prime-boost regimen and in an Ad5 seropositive host.

## Discussion

This study provides a side-by-side comparison of Ad5, mRNA, and protein vaccines across multiple immunological parameters and in different serological contexts. While previous studies, including those comparing mRNA, Ad26, and protein platforms in human volunteers, have offered important insights ^16^, their interpretation has been confounded by host variability, including wide genetic diversity and uncontrolled pre-existing immunity to adenoviruses. In contrast, our study leverages a controlled murine model to systematically assess the immunological differences between vaccine platforms, enabling clear attribution of observed effects to the vaccines themselves rather than to host background, immune histories, or genetic variation. By incorporating both naïve and Ad5-seropositive mice, we further model clinically relevant contexts that may influence vaccine efficacy.

We show that while Ad5 elicits robust immunity after a single dose in naive hosts, its relative performance declines after re-administration. In contrast, mRNA vaccines induce modest immune responses after a single dose, but robust immune responses after a booster dose. These findings underscore that vaccine performance is context dependent, shaped not only by each platform’s immunobiology, but also by the number of shots administered and the host’s serological status. It is important to note that the immunogenicity of mRNA vaccines is not entirely impervious to pre-existing immunity. In a previous study, we showed that antibody responses against the encoded antigen can accelerate clearance of this cognate antigen at the immunization site, affecting the priming of immune responses ^17^.

Our studies do not necessarily conclude that adenovirus vaccines are less effective than mRNA vaccines. There are advantages of adenoviral vaccines over mRNA vaccines, including their low cost and stability, which makes them better suited for low-income countries; as well as their potent immunogenicity, immune durability, and protective efficacy after just a single dose ^18^. Adenovirus vectors also induce more durable antigen expression lasting more than a week, potentially making them more appropriate for applications that require long-term protein expression. mRNA vaccines, on the other hand, induce an acute “burst” in antigen expression at significantly higher levels than the other vaccines as measured by in vivo bioluminescence, which could make them more appropriate for applications that require transient antigen expression. These distinct antigen kinetics likely contribute to the differences in immune responses between the vaccines. The “burst” in antigen expression observed with mRNA vaccines is associated with more potent DC activation, including higher MHC class I and costimulatory molecule expression relative to the other vaccines, which may contribute to the high anamnestic capacity of memory CD8 T cells in mRNA-immunized mice. Additionally, mRNA vaccines induce higher levels of inflammatory cytokines such as IP-10, IFN-β, and IFNγ, consistent with a robust interferon signature, which was validated at the gene expression level and the protein level. Prior studies have shown that IL-6 is critical for the adjuvant activity of LNPs ^19^, and consistent with this, we observed significantly higher IL-6 levels following mRNA vaccination compared to the other vaccine platforms, suggesting that IL-6 plays a key role in shaping immune responses to mRNA vaccines.

The protective efficacy of the three vaccines was then compared using a stringent Listeria challenge model. This challenge model showed that an mRNA prime-boost regimen confers superior protection to the other vaccine regimens. This high level of immune protection against Listeria in mRNA vaccinated mice could be explained mostly by memory CD8 T cells, given that these cells play a critical role in the clearance of intracellular infections.

Our study has limitations. First, our bioluminescence experiments rely on a luciferase reporter to quantify antigen expression. Lack of bioluminescence does not necessarily indicate complete antigen clearance, but rather the loss of luciferase’s enzymatic activity. Antigen fragments may persist beyond the detection window and continue priming immune responses. Second, we did not assess the effects of additional booster doses beyond a prime-boost regimen. Given that many individuals have already received multiple boosters during the COVID-19 pandemic, it would be valuable to investigate how repeated boosters influence immune responses across different vaccine platforms. Overall, our findings demonstrate that the relative superiority of different vaccine platforms is context dependent. After a single dose, Ad5 vaccines induce long-term antigen expression associated with robust and durable immune responses, making them well-suited as single-dose vaccination strategies. While mRNA vaccines induce modest primary CD8 T cell responses after a single dose, these responses become highly anamnestic upon boosting immunization. mRNA vaccines are also not affected by anti-vector immunity, enabling repeated use of this vaccine platform. These data highlight the importance of considering population serological background and context when selecting vaccines against novel diseases.

## ACKNOWLEDGEMENTS

Research reported in this publication was in part supported by the National Institute of Drug Abuse (NIDA) of the National Institutes of Health under the following grants to PPM: Director’s Pioneer Award (NIDA DP1DA063166). The content is solely the responsibility of the authors and does not necessarily represent the official views of the NIH. This manuscript is subject to the NIH Public Access Policy. Through acceptance of this federal funding, NIH has been given a right to make this manuscript publicly available in PubMed Central upon the Official Date of Publication, as defined by NIH.

## AUTHOR CONTRIBUTIONS

P.P.M., B.A., and S.S. designed the experiments. B.A., S.L., S.S., T.D., M. H. L., N.I., G.A. and L.A. performed the experiments. M.A. and S.M. analyzed the gene expression data. P.P.M. and B.A. wrote the manuscript with feedback from all authors.

## MATERIALS AND METHODS

### Mice and immunizations

Sex as a biological variable: Our study examined male and female animals, and similar findings are reported for both sexes. 6-8-week-old C57BL/6 mice were used. Mice were immunized intramuscularly with each respective vaccine. Mice were purchased from Jackson laboratories (approximately half males and half females) and were housed at Northwestern University’s Center for Comparative Medicine (CCM). All mouse experiments were performed with the approval of the Northwestern University Institutional Animal Care and Use Committee (IACUC).

### Reagents, flow cytometry, and equipment

Dead cells were gated out using Live/Dead fixable dead cell stain (Invitrogen). SARS-CoV-2 spike peptide pools were used for intracellular cytokine staining (ICS) and these were obtained from BEI Resources. Biotinylated MHC class I monomers were used for detecting antigen-specific CD8 T cells and were obtained from the NIH tetramer facility at Emory University. MHC class II tetramers were used for detecting antigen-specific CD4 T cells and were also obtained from the NIH tetramer facility at Emory University. Cells from PBMCs and tissues were stained with fluorescently-labeled antibodies against CD8α (53-6.7 on PerCP-Cy5.5), CD44 (IM7 on FITC), CD127 (A7R34 on Pacific Blue), KLRG-1 (2F1 on PE-Cyanine7), IL-2 (JES6-5H4 on PE), TNFα (MP6-XT22 on PE-Cy7), IFNγ (XMG1.2 on APC) or tetramers (APC or PE). Fluorescently-labeled antibodies were purchased from BD Pharmingen, except for anti-CD127 and anti-CD44 (which were purchased from Biolegend). Flow cytometry samples were acquired with a Becton Dickinson Canto II or an LSRII and analyzed using FlowJo v10 (Treestar).

### SARS-CoV-2 spike, OVA, and Ad5 hexon-specific ELISA

Binding antibody titers were measured using ELISA as described previously ^14,15,17,20–23^. In brief, 96-well flat bottom plates MaxiSorp (Thermo Scientific) were coated with 0.1μ/well of the respective protein, for 48 hr at 4°C. Plates were washed with PBS + 0.05% Tween-20. Blocking was performed for 4 hr at room temperature with 200 μL of PBS + 0.05% Tween-20 + bovine serum albumin. 6μL of sera were added to 144 μL of blocking solution in the first column of the plate, 1:3 serial dilutions were performed until row 12 for each sample, and plates were incubated for 60 minutes at room temperature. Plates were washed three times followed by the addition of goat anti-mouse or anti-human IgG horseradish peroxidase-conjugated (Southern Biotech) diluted in blocking solution (1:5000), at 100 μL/well and incubated for 60 minutes at room temperature. Plates were washed three times and 100 μL /well of Sure Blue substrate (Sera Care) was added for approximately 8 minutes. The reaction was stopped using 100 μL/well of KPL TMB stop solution (Sera Care). Absorbance was measured at 450 nm using a Spectramax Plus 384 (Molecular Devices). SARS-CoV-2 spike protein was produced in-house using a plasmid produced under HHSN272201400008C and obtained from BEI Resources, NIAID, NIH: vector pCAGGS containing the SARS-related coronavirus 2; Wuhan-Hu-1 spike glycoprotein gene (soluble, stabilized). OVA protein was purchased from Worthington Biochemical (catalog LS003049). Ad5 hexon protein was purchased from BioRad (catalog MPP002).

### Ad5, mRNA, and protein vaccines

Ad5 (10^8^ VP), mRNA (5 μg), and protein (10 μg) vaccines were diluted in PBS and administered intramuscularly (50 μL/quadriceps). These doses are standard for each vaccine and were selected based on prior dose-escalation studies showing that they elicit maximal immune responses (immune responses plateau at these doses and are not significantly improved with higher doses) ^6,24,25^. Protein vaccines were formulated 1:10 in AdjuPhos. Ad5 vaccines were purchased from the Iowa Vector Core. SARS-CoV-2 spike protein vaccine was produced in-house using a plasmid produced under HHSN272201400008C and obtained from BEI Resources, NIAID, NIH: vector pCAGGS containing the SARS-related coronavirus 2; Wuhan-Hu-1 spike glycoprotein gene (soluble, stabilized). OVA protein vaccine was purchased from Worthington Biochemical (catalog LS003049).

We synthesized mRNA vaccines in house. mRNA vaccines encoding for the codon-optimized proteins. Constructs were purchased from Integrated DNA Technologies (IDT) or Genscript, and contained a T7 promoter site for *in vitro* transcription of mRNA. The sequences of the 5′- and - 3′′-UTRs were identical to those used in a previous publication.^44^ All mRNAs were encapsulated into lipid nanoparticles using the NanoAssemblr Benchtop system (Precision NanoSystems) and confirmed to have similar encapsulation efficiency (∼95%). mRNA was diluted in Formulation Buffer (Catalog # NWW0043, Precision NanoSystems) to 0.17 mg/mL and then run through a laminar flow cartridge with GenVoy ILM encapsulation lipids (Catalog # NWW0041, Precision NanoSystems) with N/P (Lipid mix/mRNA ratio of 4) at a flow ratio of 3:1 (RNA: GenVoy-ILM), with a total flow rate of 12 mL/min, to produce mRNA–lipid nanoparticles (mRNA-LNPs). mRNA-LNPs were evaluated for encapsulation efficiency and mRNA concentration using RiboGreen assay using the Quant-iT RiboGreen RNA Assay Kit (Catalog #R11490, Invitrogen, Thermo Fisher Scientific).

### scRNA-Seq Data Acquisition

C57BL/6 mice were immunized intramuscularly with each respective vaccine. At day 1 post-vaccination, lymph nodes from five mice were pooled and MACS-sorted with a CD45 MACS positive selection kit (STEMCELL) and used for single-cell sequencing. The sorted cells were used for single-cell sequencing, and libraries were generated using 10x Genomics 3′ kits. Single-cell RNA sequencing data from mouse samples were processed using the Cell Ranger (cloud analysis) pipeline developed by 10x Genomics. Raw base call (BCL) files were demultiplexed into FASTQ files using the cellranger mkfastq command, with sample sheet information specifying index assignment. If FASTQ files were already available, this step was skipped. Reads were aligned to the mouse reference genome (mm10) using the cellranger count pipeline with default parameters. The mm10 reference transcriptome provided by 10x Genomics (refdata-gex-mm10-2020-A) was used for alignment and gene annotation. During this process, Cell Ranger performed read alignment using STAR, filtered and corrected barcodes, quantified unique molecular identifiers (UMIs), and generated a gene-cell count matrix. The resulting filtered_feature_bc_matrix directory containing the gene-barcode count matrix was used for downstream analysis. Cell Ranger also provided quality control metrics and summary statistics in the web_summary.html file. scRNA-seq was performed at the Center for Virology and Vaccine Research at BIDMC.

### scRNA-Seq Data Analysis

Gene expression analysis was performed using R version 4.5.0, Bioconductor version 3.21, and Seurat version 5.3.0. Cells were filtered to retain those expressing between 300 and 5,000 genes, with fewer than 30,000 total UMI counts, and less than 5% mitochondrial gene content. Genes detected in at least three cells were retained for downstream analysis. Data normalization was performed using the LogNormalize method in Seurat, and 2,000 highly variable genes were selected using the ’vst’ method. Doublet cells were identified and removed using the scDblFinder package (v1.10.0) with default parameters. Following normalization and scaling, principal component analysis (PCA) was performed, and samples were integrated using Seurat’s IntegrateLayers function with canonical correlation analysis (CCA) integration. Clustering was conducted using 25 principal components for both FindNeighbors and RunUMAP, and clusters were identified with a resolution of 3 using FindClusters. Cell types were annotated using the R package SingleR version 2.10.0 with the Immunological Genome Project (ImmGen) reference and refined manually based on canonical marker gene expression. The presence of Itgax (Cd11c) in clusters was used for dendritic cell annotation. For downstream analyses, cell types of interest were subset and re-normalized using the same workflow.

Differential gene expression (DEG) analysis was performed using Seurat’s FindMarkers function (Wilcoxon test), comparing each vaccine group to the naïve group, with a minimum detection threshold of 10% of cells (min.pct = 0.1). Ribosomal and mitochondrial genes were excluded from DEG results prior to downstream analysis. Genes with an absolute log2 fold change greater than 0.5 and a Benjamini-Hochberg adjusted FDR less than 0.05 were considered significant.

For pathway enrichment analysis, the fgsea package (v1.34.0) was used. Differentially expressed genes for each cell type were identified using a minimum detection threshold of 5% of cells (min.pct = 0.05). Genes were then ranked for each comparison using the metric sign(log2FC) × - log10(p-value). Gene sets included the Hallmark and C2 curated collections from MSigDB (v2023.1), the Blood Transcriptome Modules (BTM), and in-house curated gene sets. Pathways with FDR-adjusted p-values less than 0.05 were considered significantly enriched. Figures were generated using Seurat and ggplot2 (v3.5.2). Volcano plots were created using EnhancedVolcano (v1.27.0), and Venn diagrams were generated with ggVennDiagram (v1.5.2). Refer to the following papers for additional information: Seurat ^26–30^; scDblFinder ^31^; MSigDB and GSEA ^32^; R and Bioconductor ^33^

### Multiplex cytokine/chemokine assay

Blood samples were centrifugated at 15000 rpm for 10 min at 4°C to separate the serum. The serum samples were collected and frozen in -80°C until its use. A multiplex cytokines/chemokines kit was purchased from Mesoscale Diagnostics LLC or Sigma Aldrich (Luminex) and used for quantifying serum cytokines/chemokines.

### Listeria quantification

Spleens were collected from infected mice on day 3 post-challenge. Bacterial titers were quantified by homogenizing tissues through a 70 μm strainer and resuspended in 1% Triton. 10-fold serial dilutions were created in 1% Triton and added dropwise onto 6-well BHK agar plates. Plates were manually rocked and then incubated at 37°C 5% CO_2_ for 48 h. Colonies were counted the next day.

### In vivo bioluminescence

After vaccinating mice with Ad5 or mRNA expressing a luciferase reporter (Ad5-Luc or mRNA-Luc), or luciferase protein itself, luciferin (GoldBio, Catalog # LUCK-100) was administered intraperitoneally 15 min before imaging to quantify luciferase expression, as done previously ^3,17^. Mice were anesthetized and imaged using a SII Lago IVIS Imager (Spectral Instruments Imaging). Region of interest (ROI) bioluminescence was used to quantify signal.

### Statistics

Statistical tests used are indicated on each figure legend. Horizontal dashed lines in data figures represent the limit of detection. Vertical dashed lines indicate time of boosting immunizations. Data were analyzed using Prism version 10 (Graphpad).

### Study approval

#### Mouse studies

All mouse experiments were performed with approval from the IACUC of Northwestern University (study approval numbers IS00003258, IS00015002, and IS00008785). All mouse experiments with BL2 agents were performed with approval from the IACUC of Northwestern University.

#### Human specimens

De-identified human serum samples were obtained through Innovative Research Inc (samples I-598-through I-607); or through BEI Resources (samples B-655 through B-659, and 261, 012, 727, 846); these serum samples were part of other studies (secondary research).

#### Data availability

scRNA-seq data were deposited in the NCBI Gene Expression Omnibus (GEO) database under accession number GSE302672, at https://www.ncbi.nlm.nih.gov/geo/query/acc.cgi?acc=GSE302672. Other data are available upon request.

## Competing Interests

P.P.M is a consultant for Thyreos Vaccines. The other authors declare no competing interests.

---

**Supplemental Figure 1.**
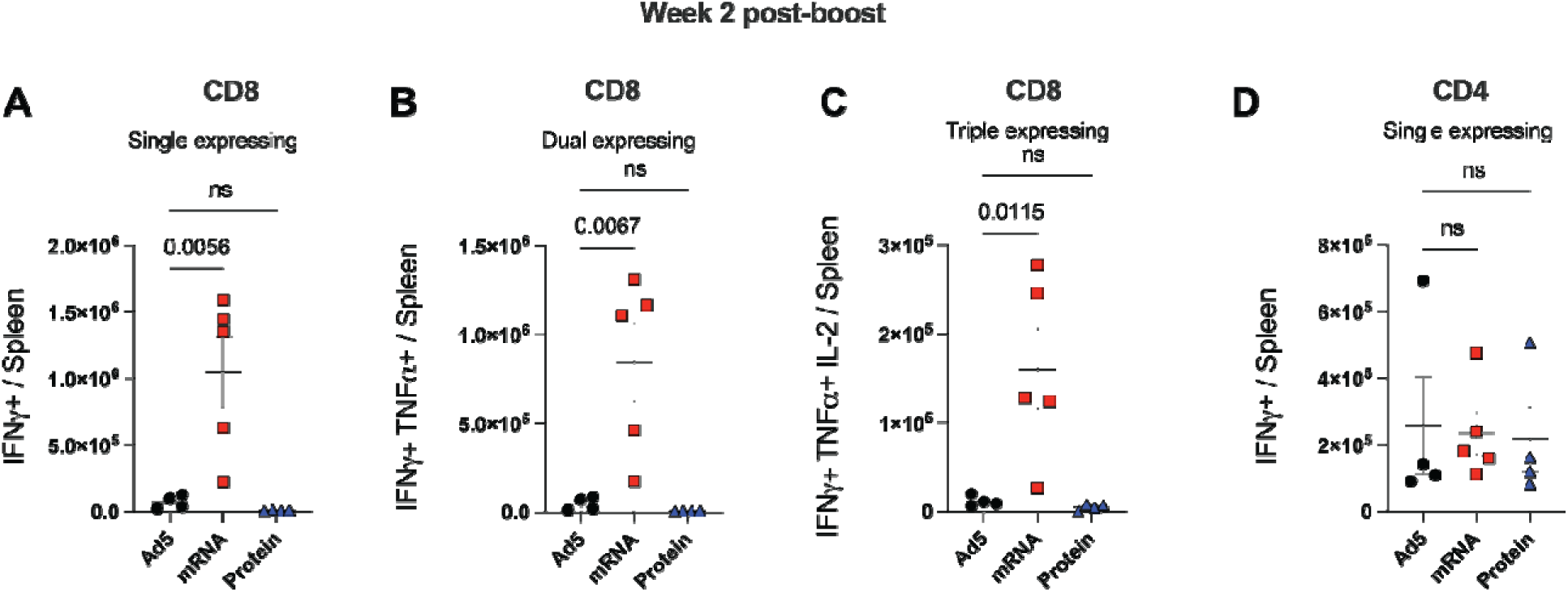
Cytokine expression by virus-specific T cells following immunization with Ad5, mRNA, and protein vaccines. Experimental outline was similar to that in Figure 6A. Numbers of spike-specific CD8 T cells that express IFN-*_γ_* (**A**); IFN-*_γ_* and TNF-_α_ (**B**); and IFN-*_γ_*, TNF-IL-2 (**C**) in spleen at week 2 post-boost. (**D**) Numbers of spike-specific CD4 T cells that express IFN-*_γ_* in spleen at week 2 post-boost. Spike peptide pool stimulations were performed for 5 h. Data from one experiment (n=5 mice per group). Experiment was repeated once with similar results. Indicated *P* values were calculated by ordinary one-way ANOVA with Dunnett’s multiple comparisons. Error bars represent SEM.

**Supplemental Figure 2.**
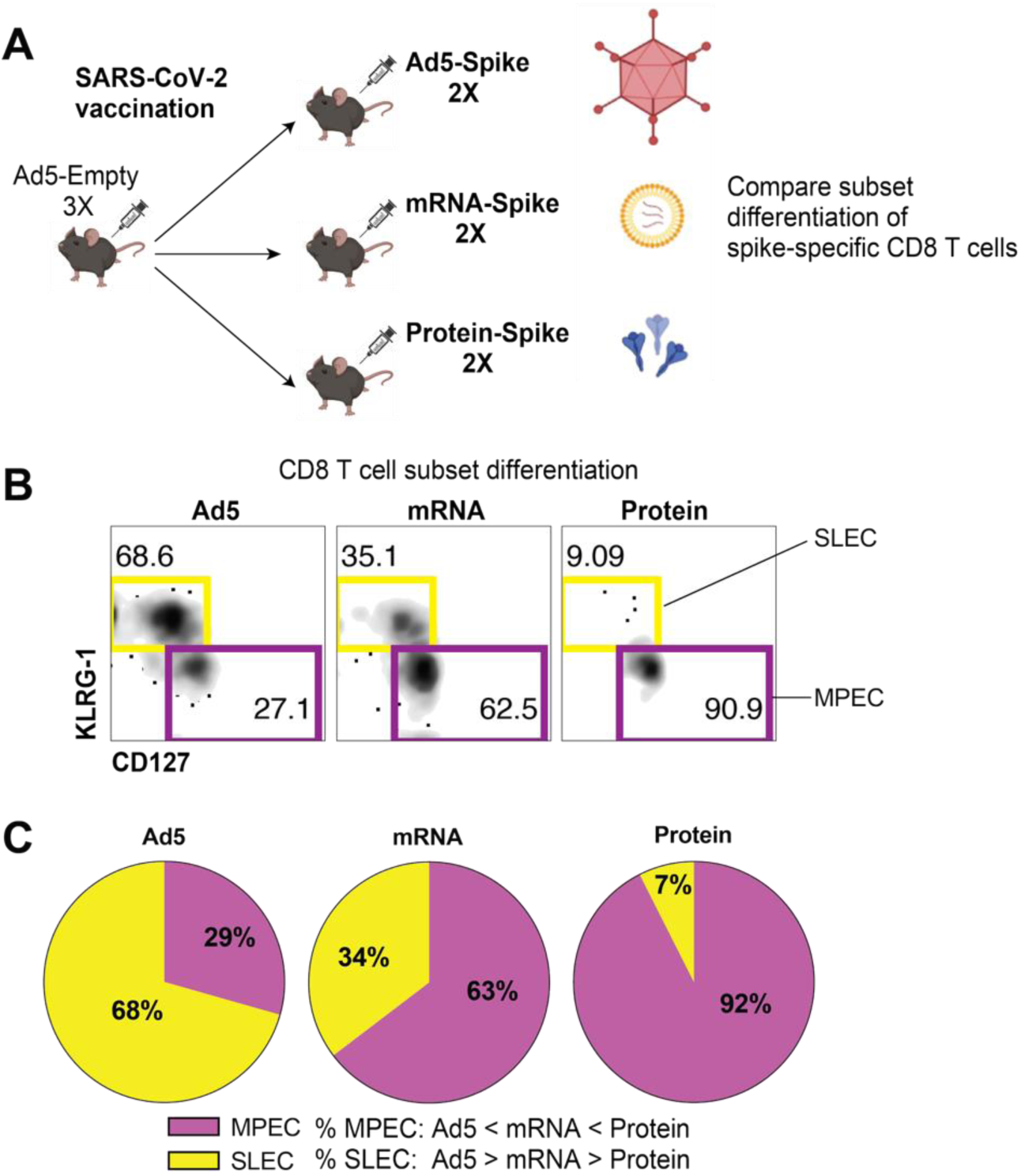
Comparative analyses of CD8 T cell subset differentiation. (**A**) Experimental outline wa similar to that in Figure 6A. (**B**) Representative FACS plots of short-lived effector cells (SLECs) and memory precursor effector cells (MPECs) populations, gated on SARS-CoV-2–specific (K^b^ VL8+) CD8 T cells (**C**) Pie diagrams showing CD8 T cell subsets. Data from one experiment (n=4-5 mice per group). Experiment was repeated once with similar results.

**Supplemental Figure 3.**
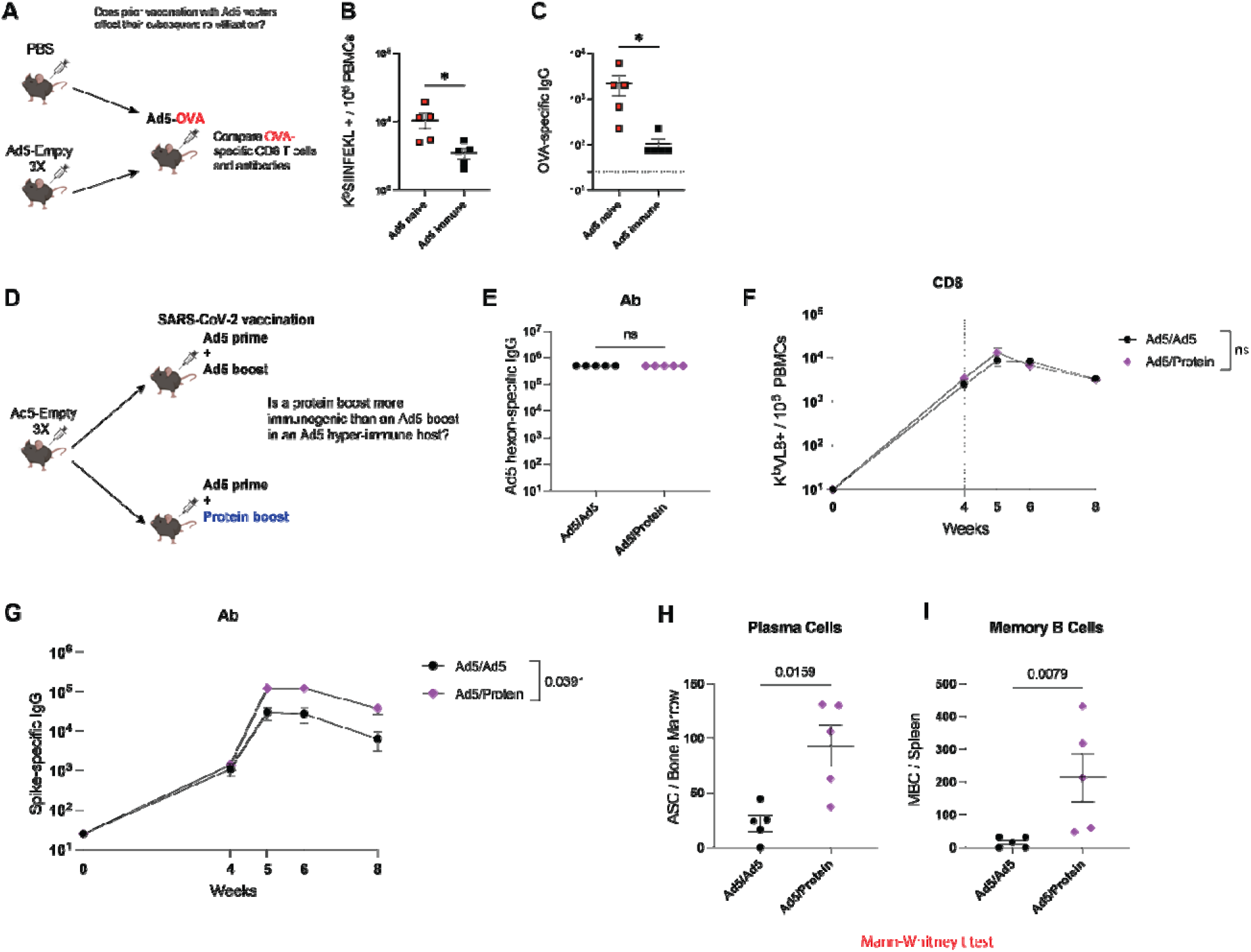
Prior immunization with Ad5 renders Ad5 vectors les immunogenic. (**A**) Experimental outline for comparing Ad5 vaccine-elicited responses in mice that had been previously immunized with Ad5 vectors. First, Ad5 seropositivity was induced by injecting C57BL/6 mice with Ad5-Empty, injected intramuscularly, once every 3 weeks for a total of three doses. Control mice were injected with PBS. Mice were then immunized with Ad5-OVA and immune responses were measured at week 2 post-immunization. (**B**) Summary of OVA-specific (K^b^ SIINFEKL+) CD8^+^ T cells in PBMCs. (**C**) Summary of OVA-specific antibody titers in sera. **(D**) Experimental outline for determining whether a heterologou Ad5/protein regimen elicits superior immune responses, relative to a homologous Ad5/Ad5 regimen. (**E**) Summary of Ad5 hexon-specific antibody responses in sera. (**F**) Summary of spike-specific CD8 T cells in PBMCs. (**G**) Summary of spike-specific antibody responses in sera. (**H**) Spike-specific plasma cell responses in bone marrow. (**I**) Spike-specific memory B cell responses in spleen. Data from one experiment (n=5 mice per group). Experiment was repeated once with similar results. In panels B,C, E, and F indicated *P* values were calculated by Mann-Whitney t test (p value from panel F is from week 8). In panel G, indicated *P* value was calculated by Welch’s t test. Error bars represent SEM.

**Supplemental Figure 4.**
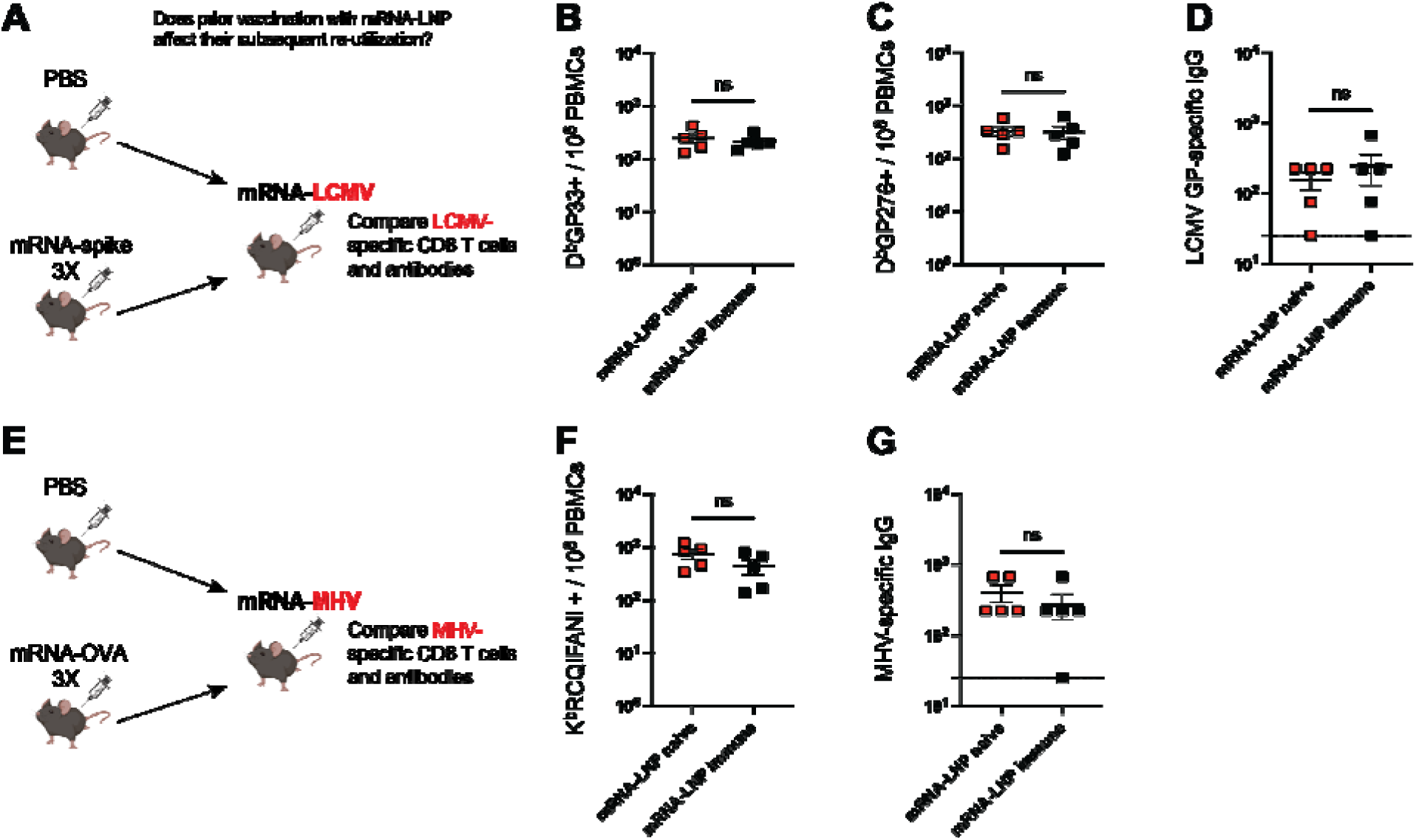
Prior immunization with mRNA vaccines does not impair thei subsequent re-utilization. (**A**) Experimental outline for comparing mRNA vaccine-elicited responses in mice that had been previously immunized with mRNA-spike vaccines. (**B**) Summary of LCMV GP33-specific CD8^+^ T cell responses in PBMCs. (**C**) Summary of LCMV GP276-specific CD8^+^ T cell responses in PBMCs. (**D**) Summary of LCMV GP-specific antibody responses in sera. (**E**) Experimental outline for comparing mRNA vaccine-elicited responses in mice that had been previously immunized with mRNA-OVA vaccines. (**F**) Summary of MHV-specific CD8^+^ T cell responses (K^b^RCQIFANI+) in PBMCs. (**G**) Summary of MHV-specific antibody responses in sera. Data from panels B, C, D, F, and G are from week 2 post-immunization. Data from one experiment (n=5 mice per group). Indicated *P* values were determined by Mann-Whitney t test. Error bars represent SEM.

**Supplemental Figure 5.**
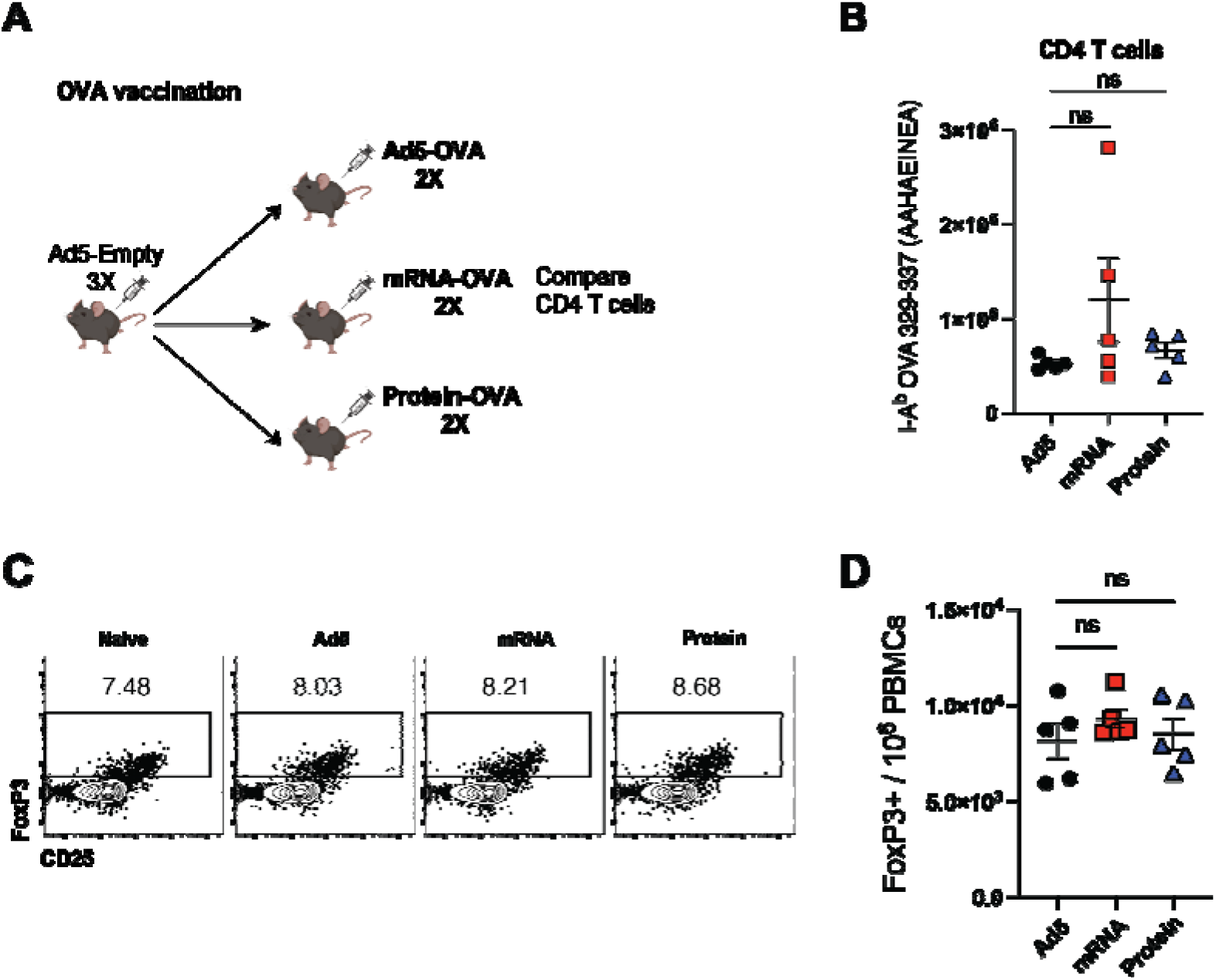
Similar CD4 T cell responses among different vaccine platforms. (**A**) Experimental outline similar to Fig. 7A. (**B**) Summary of OVA-II–specific (I-A^b^ AAHAEINEA+) CD4^+^ T cells in the spleen at week 8. (**C**) Representative FACS plots showing FoxP3+ CD4+ T regulatory cells in PBMCs at week 2. (**D**) Summary of Treg responses in PBMCs at week 2. Data from one experiment (n=5 mice per group). Experiment was repeated once with similar results. Indicated *P* values were calculated by ordinary one-way ANOVA with Dunnett’s multiple comparisons. Error bars represent SEM.

## Notes

### Competing Interest Statement

The authors have declared no competing interest.

